# Synaptotagmin-1 and Doc2b exhibit distinct membrane remodeling mechanisms

**DOI:** 10.1101/538405

**Authors:** Raya Sorkin, Margherita Marchetti, Emma Logtenberg, Melissa Piontek, Emma Kerklingh, Guy Brand, Rashmi Voleti, Josep Rizo, Wouter H. Roos, Alexander J. Groffen, Gijs J. L. Wuite

## Abstract

While the role of Synaptotagmin-1 in living cells has been described in detail, it remains a challenge to dissect the contribution of membrane remodelling by its two cytoplasmic C2 domains (C_2_AB) to the Ca^2+^-secretion coupling mechanism. Here, we study membrane remodeling using pairs of optically-trapped beads coated with SNARE-free synthetic membranes. We find that the soluble C_2_AB domain of Syt1 strongly affects the probability and strength of membrane-membrane interactions in a strictly Ca^2+^- and protein-dependent manner. A lipid mixing assay with confocal imaging reveals that at low Syt1 concentrations, no hemifusion is observed. Notably, for similar low concentrations of Doc2b hemifusion does occur. Consistently, both C_2_AB fragments cause a reduction in the membrane bending modulus, as measured by an AFM-based method. This lowering of the energy required for membrane deformation likely contributes to the overall Ca^2+^-secretion triggering mechanism by calcium sensor proteins. When comparing symmetrical (both sides) and asymmetrical (one side) presence of protein on the membranes, Syt1 favors an asymmetrical but Doc2b a symmetrical configuration, as inferred from higher tether probabilities and break forces. This provides support for the direct bridging hypothesis for Syt-1, while hinting to possible preference for protein-protein (and not protein-membrane) interactions for Doc2b. Overall, our study sheds new light on the mechanism of Ca^2+^ induced fusion triggering, which is essential for fundamental understanding of secretion of neurotransmitters and endocrine substances.

## Introduction

Ca^2+^ sensor proteins tightly control the secretion of neurotransmitters and endocrine substances. After the arrival of an action potential to the synaptic terminal, Ca^2+^ influx triggers the fast fusion of synaptic vesicles with the presynaptic plasma membrane. This process depends on SNAREs, Munc18-1, Munc13s, complexins and Ca^2+^ sensor proteins such as Synaptotagmin-1 (Syt1), among other proteins (1). Syt1 contains a single transmembrane domain and two cytoplasmic C2 domains (together named C_2_AB). The C_2_A and C_2_B domains can bind three and two Ca^2+^ ions, respectively, accompanied by binding to phosphatidylserine in the membrane (2). The C_2_AB domain also binds to phosphatidylinositol 4,5-bisphosphate and SNARE proteins(3–6). Ca^2+^ binding is thought to be the direct trigger for vesicle fusion (1, 2, 7). Other C_2_-domain-containing proteins such as Doc2b, which contributes to spontaneous neurotransmitter release, similarly interact with membranes and SNARE proteins to trigger fusion(8, 9).

Despite extensive research, the exact role of Syt1 in membrane fusion, and in particular, its membrane-bound configuration before and during fusion, remain uncertain. Several mechanisms have been suggested; the “clamping hypothesis” proposes that Syt1 (together with complexin) prevents fusion in the absence of Ca^2+^ signal(10). Syt1 is also proposed to bring the vesicle and plasma membranes into close proximity upon Ca^2+^ binding, thus assisting fusion (11, 12). The C_2_AB domain of Syt1 was demonstrated to directly bridge the membranes through simultaneous binding of the Ca^2+^-binding loops and other regions of the C_2_B domain – coined the ‘direct bridging mechanism’(12, 13). These other regions may involve conserved basic lysines(14–16) (K326,327) and, although debated, two conserved arginines(12, 17) (R398 and R399). It was also demonstrated that upon Ca^2+^ binding, the Ca^2+^-binding loops not only associate with, but insert into the membrane (18–20). This insertion penetrates one leaflet of the membrane to approximately the depth of lipid glycerol backbones (7, 21), thereby inducing local membrane curvature (22–24). Full membrane fusion is associated with complete SNARE complex assembly, leading to pore formation and release of the vesicle cargo, while Syt1 contributes also to pore expansion (25–27). Two other studies led to another possible mechanism involving the interaction of protein oligomers bound on both membranes or so called ‘oligomerization mechanism’ (28, 29), although such oligomerization may arise because of insufficient purification of the soluble synaptotagmin-1 fragment used (13). Nevertheless, very recent work proposed a different mechanism according to which synaptotagmins form ring-like oligomers that facilitate vesicle docking, while their disassembly, coupled to Ca^2+^ influx, triggers SNARE-driven fusion (30–32). There is also evidence that Syt1-SNARE interactions are critical for neurotransmitter release, although the relevant binding mode(s) are still under debate (3–5). In summary, despite the wealth of information available, the mechanisms by which Syt1 and other Ca^2+^ sensor proteins facilitate membrane fusion remain unclear.

Here, we study membrane remodeling using pairs of optically-trapped beads coated with SNARE-free synthetic membranes. Through the *in vitro* observation of single membrane-membrane interactions we aim to unravel how direct C_2_AB-phospholipid interactions contribute to initiation of membrane fusion. We reveal distinct differences in membrane interactions of two structurally similar calcium sensor proteins, providing us with a mechanistic understanding why Doc2b and Syt1 act differently. Moreover, the results support the idea that Syt1 acts according to a direct bridging mechanism and Doc2b might have preference for protein-protein interactions. Overall, our study provides new insights into the role of calcium sensor proteins in membrane fusion, and can be readily extended to explore other membrane fusion events.

## Results

### Membrane interactions induced by Syt1-C_2_AB are Ca^2+−^ and protein-concentration dependent

We probed membrane-membrane interactions induced by Syt1-C_2_AB using a combination of optical tweezers and confocal fluorescence microscopy. Two microspheres (3.84 μm in diameter) coated with single phospholipid bilayers (PC:PS:Chol, 50:20:30, unless otherwise specified) were brought into contact by an automated approach-and-separation method, as previously described (33) (Fig. 1a). By monitoring the forces upon approach and retraction of the beads in the presence of proteins, we can: i) detect the presence of tethers from the change in force (see event 3 in Fig. 1a and force-time plot in Fig, 1b); ii) quantify the tether strength, which is the force at which the beads become disconnected upon membrane pulling (rupture force, Fig. 1b); iii) visualize, using confocal microscopy, single tethers in the presence of labelled proteins (Syt1-C_2_AB-mCherry, Fig.1c top panel) or labelled phospholipids (1% Rhodamine-PE, Fig. 1c bottom panel).

**Figure 1.**
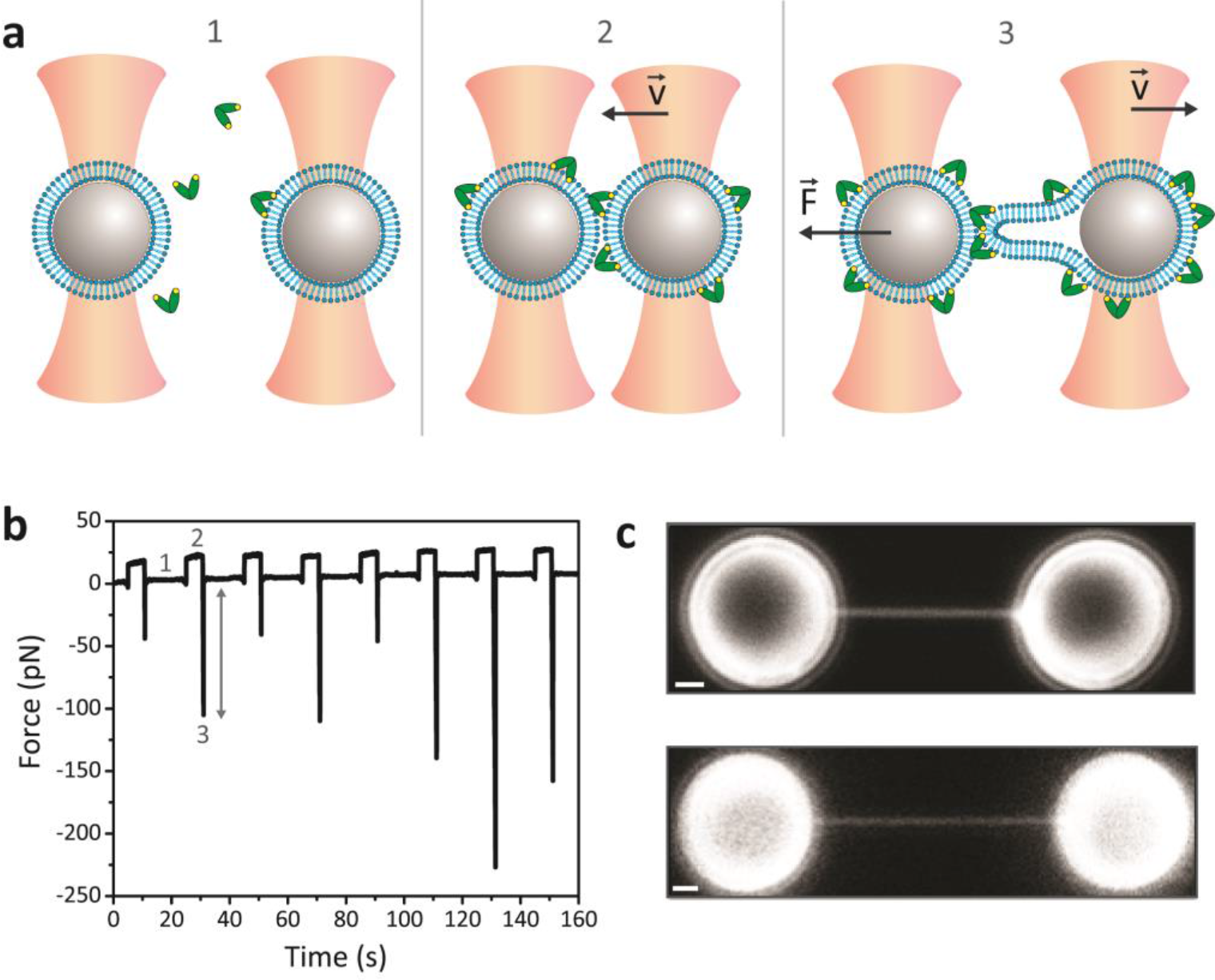
Membrane interactions captured with optical tweezers and confocal fluorescence microscopy. (a) Schematic (not to scale) of the dual-beam optical trapping setup used to manipulate two polystyrene microbeads (grey) coated with a phospholipid bilayer (blue) in the presence of proteins (green) and Ca^2+^ ions (yellow dots), panel 1. The liposomes are brought into contact for 5 s (panel 2) and moved away (panel 3) with a constant velocity, by an automated approach-and-separation method. The force on the left fixed trap is measured. (b) Typical force-time plot showing 8 consecutive interactions. The force is zero when the liposomes are apart (1). A positive force occurs during membrane contact (2) and a negative force occurs during bead separation (3), indicative of tether formation. The rupture force (indicated by grey arrow) is used to quantify the strength of each tether. (c) Confocal fluorescence images of membrane tethers visualized in the presence of labelled proteins (Syt1-C_2_AB-mCherry, top panel) or labelled phospholipids (1% Rhodamine-PE, bottom panel) on both beads. Scale bars: 1 μm.

To use this approach for studying C_2_AB – membrane interactions, we first established that the observed membrane tethers were Syt1-C_2_AB–dependent (Fig. 2 and supplementary Fig. S1). By increasing the Ca^2+^ concentration in the presence of 0.2 μM Syt1-C_2_AB in solution, the probability of interactions increased from 40% at 100 μM Ca^2+^ to 83% at 500 μM Ca^2+^ (Fig. 2a). In the absence of protein, the probability of tether formation was lower than 3% at any of the Ca^2+^ concentrations tested (Fig. 2a). The normalized cumulative distributions of the tether rupture forces (Fig. 2b, inset) and the median of these distributions (i.e. the forces at which half of the observed interactions break; Fig. 2b) reveal that the tether strength is approximately proportional to the Ca^2+^ concentration. Furthermore, the probability of interactions (Fig. 2c) and the median rupture forces (Fig 2d) at constant Ca^2+^ concentration (500 μM) increased as a function of the protein concentration. These results demonstrate that our method is suitable for detecting C_2_AB specific interactions.

**Figure 2.**
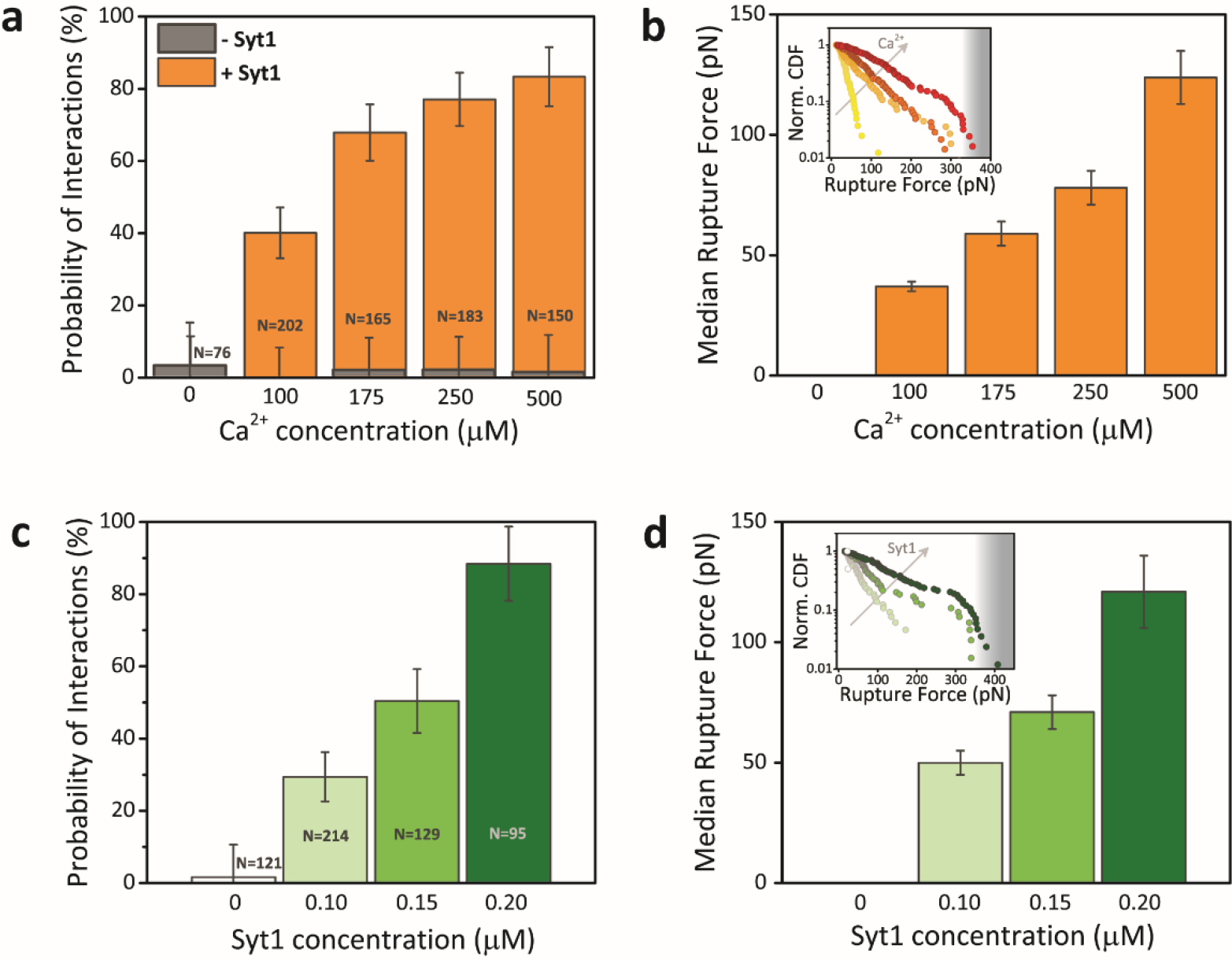
Membrane interactions induced by Syt1-C_2_AB are Ca^2+−^ and protein-concentration dependent. (a) Probability of membrane interactions with increasing Ca^2+^ concentration (in the absence of proteins or presence of 0.2 μM Syt1 C_2_AB, grey and orange histogram, respectively). Number of approach and separation cycles (N) are indicated. At least ten bead pairs were measured for each sample, with ten cycles for each bead pair. Error bars are statistical error. (b) Median tether rupture forces at different Ca^2+^ concentrations, error bars are SD from bootstrapping. Inset: Normalized cumulative distribution functions (CDF) of the rupture forces (colour coded from yellow to red as the Ca^2+^ increases), from which the median forces are calculated. (c) Probability of membrane interactions with increasing Syt1-C_2_AB concentration (at fixed 0.5 mM CaCl_2_). Number of interactions (N) are indicated. Error bars are statistical error. (d) Median rupture forces with increasing Syt1-C_2_AB concentration. Error bars are SD from bootstrapping. Inset: normalized CDF of the rupture forces recorded (colour coded from light to dark green as the Syt1 concentration increases). In (b) and (d), the grey background gradient marks the maximum force that can determined with the optical trap at the set laser power (5 W) and beads size (3.84 μm). Error bars in rupture forces plots are standard deviation of the bootstrapped median rupture force values.

### Optimal membrane bridging

To investigate how the above described membrane binding activity is supported by different protein-membrane binding modes we employed two experimental arrangements, called “symmetric” and “asymmetric”. In the asymmetric configuration, illustrated in Fig. 3a, upper panel, the proteins were bound only on one membrane-coated microsphere and brought into contact with a protein-free membrane on the other microsphere. In the symmetric case, proteins were bound to both membranes, as schematically shown in Fig. 3a (bottom panel) and visualized with fluorescence in Fig. 1c. The confocal imaging confirms that, in the asymmetric configuration, Syt1-C_2_AB-mCherry remained bound only to a single membrane-coated microsphere, suggesting that protein redistribution by membrane dissociation and re-association was minimal over the time course of the experiment. Syt1-C_2_AB-mCherry was also present on a tether that extended from the protein-coated bead (fluorescence image Fig. 3a, middle panel, and supplementary Fig. S2, a). This tether configuration, i.e. extension from one of the beads, was observed also in symmetric protein configurations with a single phospholipid-labeled bead (supplementary Fig. S2). We note that all force measurements presented in this manuscript were obtained with unlabeled proteins, while mCherry Syt-1 was only used for imaging. As shown in Fig. 3b, Syt1-C_2_AB was more effective in the asymmetric configuration, resulting in a more than 2-fold higher probability of interactions and 3-fold stronger tethers (Fig. 3b). This behavior was consistent at different Syt1 concentrations, as shown in supplementary Fig. S1, which also demonstrates that the protein was not saturated under these conditions. Our liposome aggregation assay provides further evidence that the protein concentrations used throughout the optical tweezers experiments are below the saturation limit, as shown in supplementary Fig. S3. Performing the same experiments with Doc2b revealed the opposite behavior: the probability of interactions and rupture forces were strongly enhanced in the symmetric configuration (Fig. 3c). Statistical significance of the results was determined by a two sample Kolmogorov–Smirnov test. This is a two-tailed nonparametric test that quantifies the distance between the empirical distributions of two data sets, where the null hypothesis states that the two samples are drawn from the same distribution. Rupture forces in symmetrical vs. asymmetrical configurations were found to be significantly different for both Syt-1 and Doc2b with P<0.05. The distinct results obtained with the two proteins are unlikely to arise from differences in their membrane affinities, as measurements of liposome aggregation as a function of protein concentration yielded similar EC50 values for the two proteins (Fig. S3). The activity of Syt1-C_2_AB showed an optimum at 0.8 μM in the liposome aggregation assay. The reduced clustering activity at higher Syt1 concentrations may be due to prevalence of symmetric (protein-protein) interactions under such conditions.

**Figure 3.**
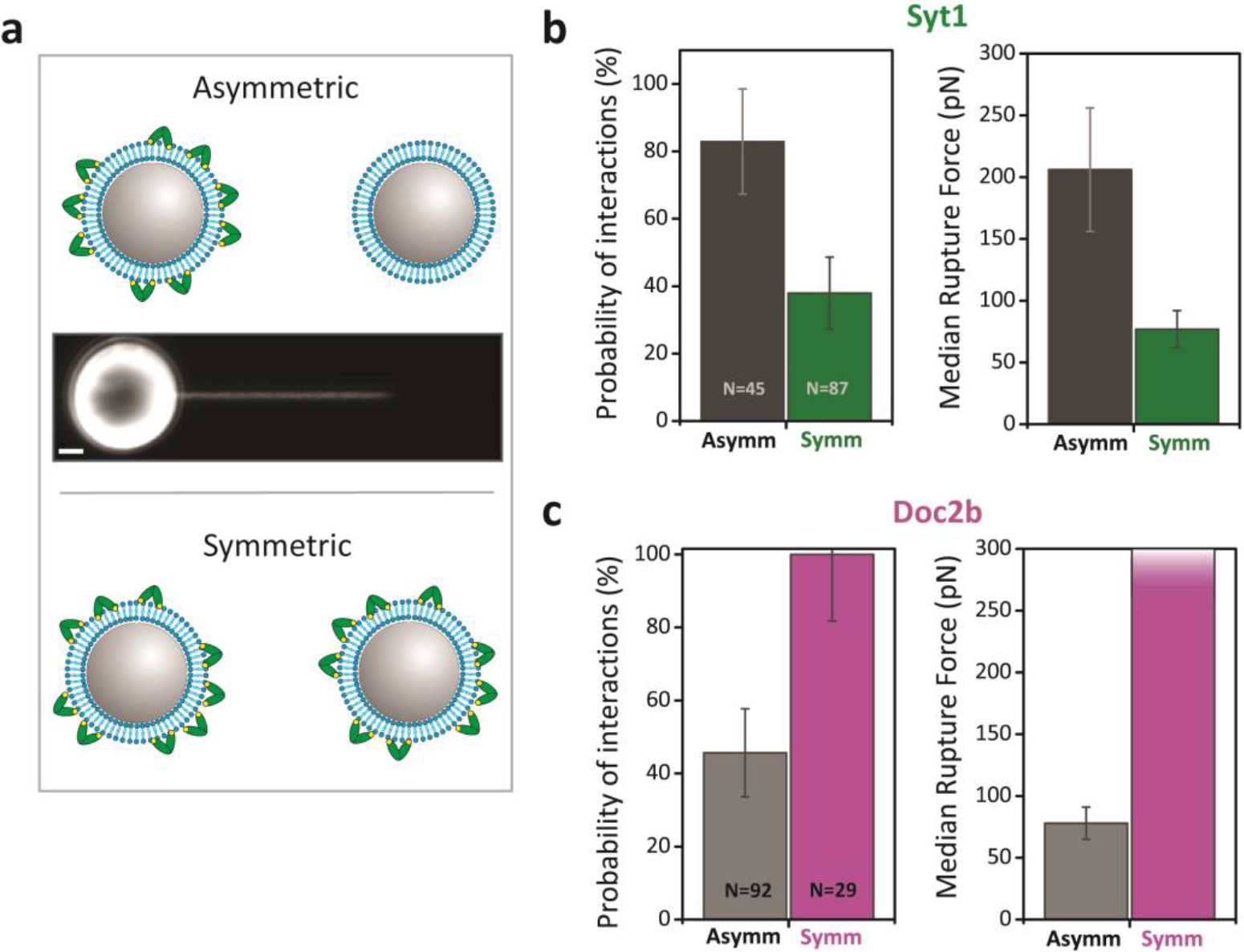
Optimal tethering by Syt1 and Doc2b loaded on single and dual membranes, respectively. (a) Schematic of the two experimental configurations used. Asymmetric: protein was bound to a single membrane-coated bead (upper panel), as confirmed by confocal imaging where Syt1-C_2_AB-mCherry (0.5 μM Syt1, 0.25 mM CaCl2) was bound to the left bead and decorated the tether structure, but not the other bead. Note that at other times the tether was drawn from the dark bead and no bias of tether extension towards labelled lipids was observed. Symmetric: proteins were bound to both beads (bottom panel, see Fig 1C for confocal image). Scale bar: 1μm. (b) Left: Probability of Syt1-mediated membrane interactions in the asymmetric (dark grey) and symmetric (green) configurations. Error bars are statistical error. Right: Median rupture force in the two configurations. Error bars are SD from bootstrapping. (c) Left: Probability of Doc2b-mediated membrane interactions in the asymmetric (light grey) and symmetric (magenta) configurations. Error bars are statistical error. Right: Median rupture force in the two configurations. The gradient in the symmetric configuration indicates a lower limit of this quantification because it reached the upper limit of our trapping force. Error bars are standard deviation of the bootstrapped median rupture force values. Note: the lower concentration of Doc2b compared with Syt1 is so chosen in order to be within the range of break forces that can be experimentally quantified.

### The impact of cholesterol

The presence of cholesterol is known to increase the efficiency of membrane fusion, likely by facilitating membrane curvature changes that lower the energy barrier for fusion (34, 35). To explore the effect of cholesterol on C_2_AB-mediated membrane-membrane interactions by Syt1 and Doc2b, we performed the same experiments in the presence or absence of cholesterol (PC:PS, 80:20 and PC:PS:chol, 50:20:30). We tested the two Ca^2+^ sensors in their most efficient configurations, (i.e. asymmetric for Syt1 and symmetric for Doc2b). The probability of interactions did not increase further for Syt1-C_2_AB (Fig. 4a, panel i), but an increase in the median rupture force was observed (Fig. 4a, right panel). In the case of Doc2b, cholesterol significantly increased both the probability of interactions and the tether strength (Fig. 4b, cumulative distribution functions for the data presented in supplementary Fig. S4). Statistical significance of the results was determined by a two sample Kolmogorov–Smirnov test. Using this test to compare the distributions of tether break forces with and without added cholesterol in the membrane, we find that the difference for Doc2b and Syt-1 mediated tether break forces are statistically significant with P<0.005.

**Figure 4.**
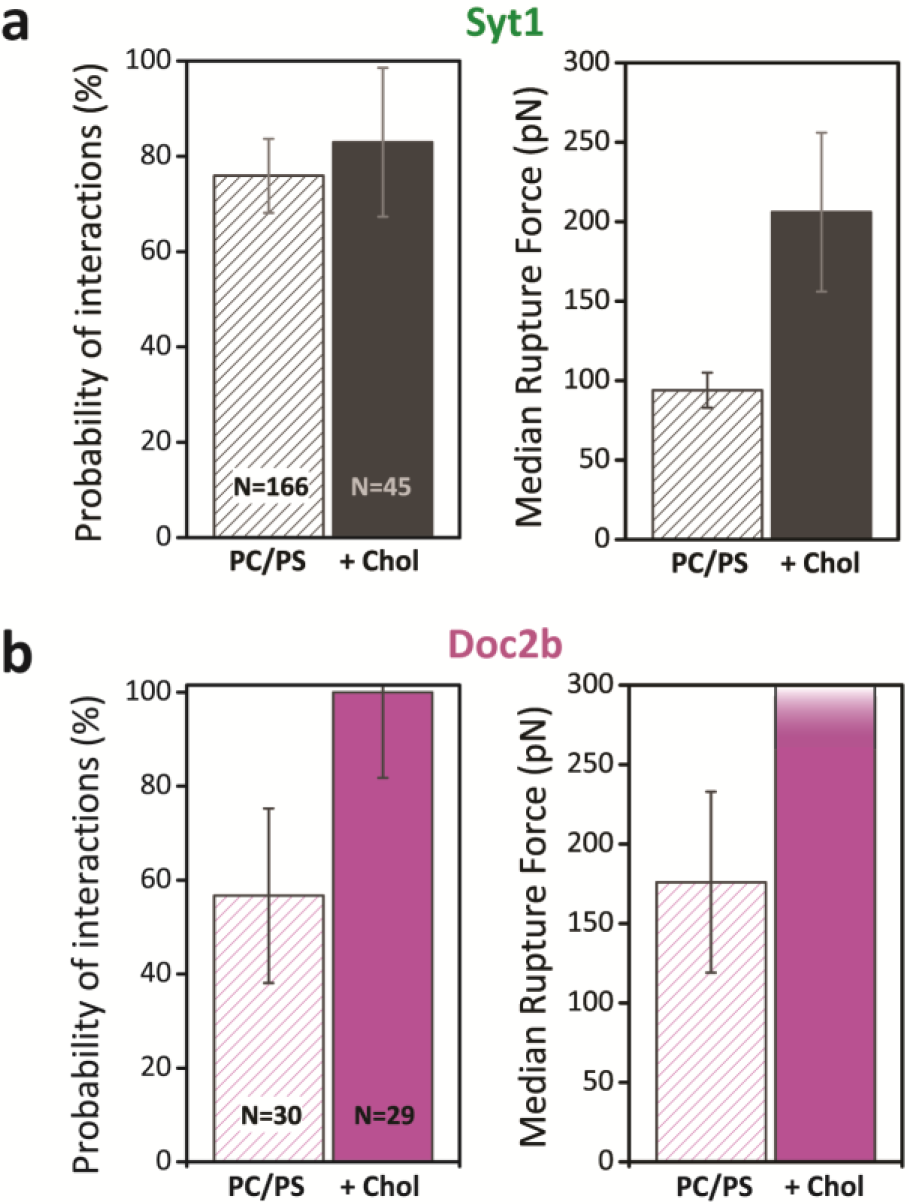
Cholesterol significantly increases strength and probability for Doc2b mediated tethers. Probability of interactions and median rupture forces for membranes containing only PC/PS (80:20) and in the presence of cholesterol (30%, + Chol). (**a**) Syt1-mediated interactions tested in the asymmetric configuration (0.5 μM, 0.25 mM CaCl_2_). (**b**) Doc2b-mediated interactions tested in the symmetric configuration (0.05 μM, 0.25 mM CaCl_2_). Number of approach and separation cycles (N) are indicated. Error bars in probability plots are statistical error; error bars in rupture forces plots are standard deviations of the bootstrapped median rupture force values.

### Syt1 and Doc2b can form different membrane tethers

To understand the membrane configuration in the tether structure, we used live confocal fluorescent microscopy to differentiate between protein-mediated membrane bridging and hemifusion events. To this end, one of the trapped beads was labeled with a fluorescent phospholipid tracer (PC:PS:Rhodamine-PE, 79:20:1 or PC:PS:chol:Rhodamine-PE, 49:20:30:1) while the other bead was unlabeled (PC:PS, 80:20 or PC:PS:chol 50:20:30). The beads were brought into contact and separated to a distance of 200-300 nm in the presence of Syt1 or Doc2b, with the whole process of approach, waiting in contact and separation to a small distance lasting ~10 sec.

Once tether formation was measured as a force increase, continuous confocal imaging was used to monitor the fluorescence of the dark bead. If hemifusion occurs, phospholipid mixing is expected to cause a fluorescence increase in the outer leaflet of the dark bead (see schematic in Fig. 5a). The same method was previously used to demonstrate that Doc2b can induce hemifusion while lack of full fusion is evident from a lipid mixing assay(33). (We cannot completely exclude the less likely possibility of lipid transfer mediated by the protein.) Typical images from such experiments are shown in Fig 5b, for one bead pair in the presence of 0.5 μM Syt1 (green) showing no hemifusion, and one bead pair in the presence of Doc2b (magenta) at 0.7 μM showing hemifusion, as seen also from the corresponding fluorescence intensity for this bead pair in Fig 5c, where the fluorescent signal of the dark bead increased markedly due to lipid mixing. This concentration, which is higher than used in Fig. 3, was experimentally chosen in order to increase the detected hemifusion events. To explore the conditions that allow hemifusion, we tested multiple bead pairs at different protein concentrations and different configurations (symmetric/asymmetric), as well as different lipid compositions (with/without cholesterol). These measurements are summarized in Fig 5d; for low concentration of Syt1 C_2_AB (0.5 μM), no hemifusion occurred regardless of the presence of cholesterol and regardless of the symmetric or asymmetric configuration. However, at high Syt1 C_2_AB concentration, we observed a remarkable effect: while no hemifusion occurred in the symmetric configuration, the asymmetric configuration yielded hemifusion in 53% of the cases. Doc2b induced hemifusion already at low concentration of 0.7 μM. These lipids-mixing experiments are limited by an experimental time window of around 10 minutes due to photobleaching. The observed intensity decrease between 300 – 700 s in Fig 5c is caused by such photobleaching. We conclude that the membrane tethers formed by Syt1-C_2_AB typically represent proteolipid structures held together by membrane bridging, unless a high concentration and asymmetric configuration are used, whereas Doc2b induced tethers can represent either membrane-bridged or hemifused states.

**Figure 5.**
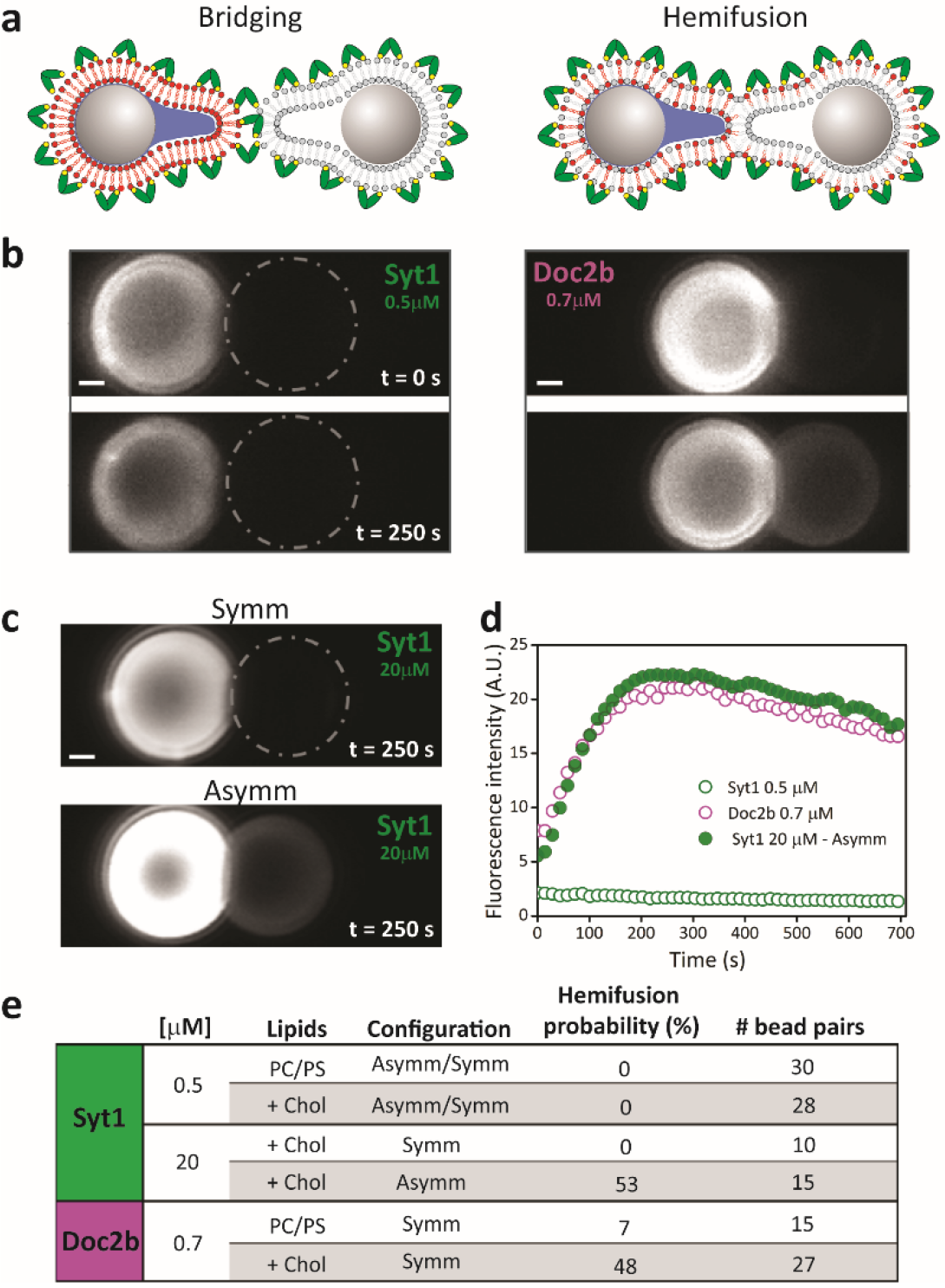
Membrane bridging and hemifusion induced by Syt1 and Doc2b. (a) Schematic of possible protein-mediated membrane interactions (proteins bound on the membrane are represented in green). Left panel: membranes bridging, where the bilayers stay separated and therefore labeled phospholipids (represented in red) remain on one membrane. Right panel: hemifusion, where the proximal membrane leaflets have fused, causing lipid-mixing. (b) Confocal images where only one membrane (left bead) was fluorescently labelled (1% Rhodamine-PE). Imaging was initiated as soon as a tether was detected. Upper panels: images recorded at t = 0 s. Lower panels: images recorded after 250 s. Observed interactions mediated by Syt1 (left panel) and by Doc2b (right panel). (c) Confocal images recorded after 250s interaction in the presence of 20 μM Syt1 in the symmetric configuration (upper panel) and in the asymmetric configuration (bottom panel). (d) Fluorescence signal recorded from the dark bead over time in the presence of 0.7 μM Doc2b and 0.5 μM or 20 μM Syt1 in (b). (e) Hemifusion probabilities under different conditions.

### Doc2b and Syt1 reduce the membrane bending modulus

Several studies demonstrated that upon Ca^2+^ binding, C_2_AB domains insert into the lipid membrane (7, 18). Such insertion is expected to induce membrane curvature (22, 23). As it disturbs the packing of the lipid acyl chains, this might result in a change in the mechanical properties of the membrane. To quantitatively study how Doc2b and Syt1 C_2_AB fragment insertion into the negatively charged membrane influences its mechanical properties, we used a recently developed method based on atomic force microscopy (AFM) (36–38). By first imaging liposomes (Fig. 6a) and then indenting them with the AFM tip, we can measure the membrane radius, stiffness and membrane tension of single liposomes. These experimentally determined parameters are then fitted to a Canham-Helfrich based model from which the bending modulus is derived (36). A phospholipid composition of 44% cholesterol, 20% egg PS, 21% egg PC and 15% Egg SM was chosen because it results in stable liposomes which do not rupture during the experiments. The extruded liposomes, with a 120±30 nm average diameter (by DLS), were found to have a bending modulus κ = 9±2 k_B_T. When performing the experiments in the presence of 0.9 μM Syt-1, κ = 9±2 k_B_T was obtained, similar to the result in the absence of protein. In the presence of a high protein concentration (20 μM Syt1-C_2_AB), which is in the same range as the concentration previously estimated by western blot analysis of isolated synaptosomes(39) and the concentration when we observe hemifusion induced by Syt1, the bending modulus was 3-fold reduced (Fig. 6c). When performing these experiments in the presence of Doc2b, a reduction in the bending modulus was observed already at 0.9 μM protein concentration. These results suggest an active role of C_2_AB domains in membrane remodeling during the fusion process: in addition to bringing the membranes into close proximity, Syt1-C_2_AB and Doc2b may also contribute to hemifusion by directly lowering the energy barrier.

**Figure 6.**
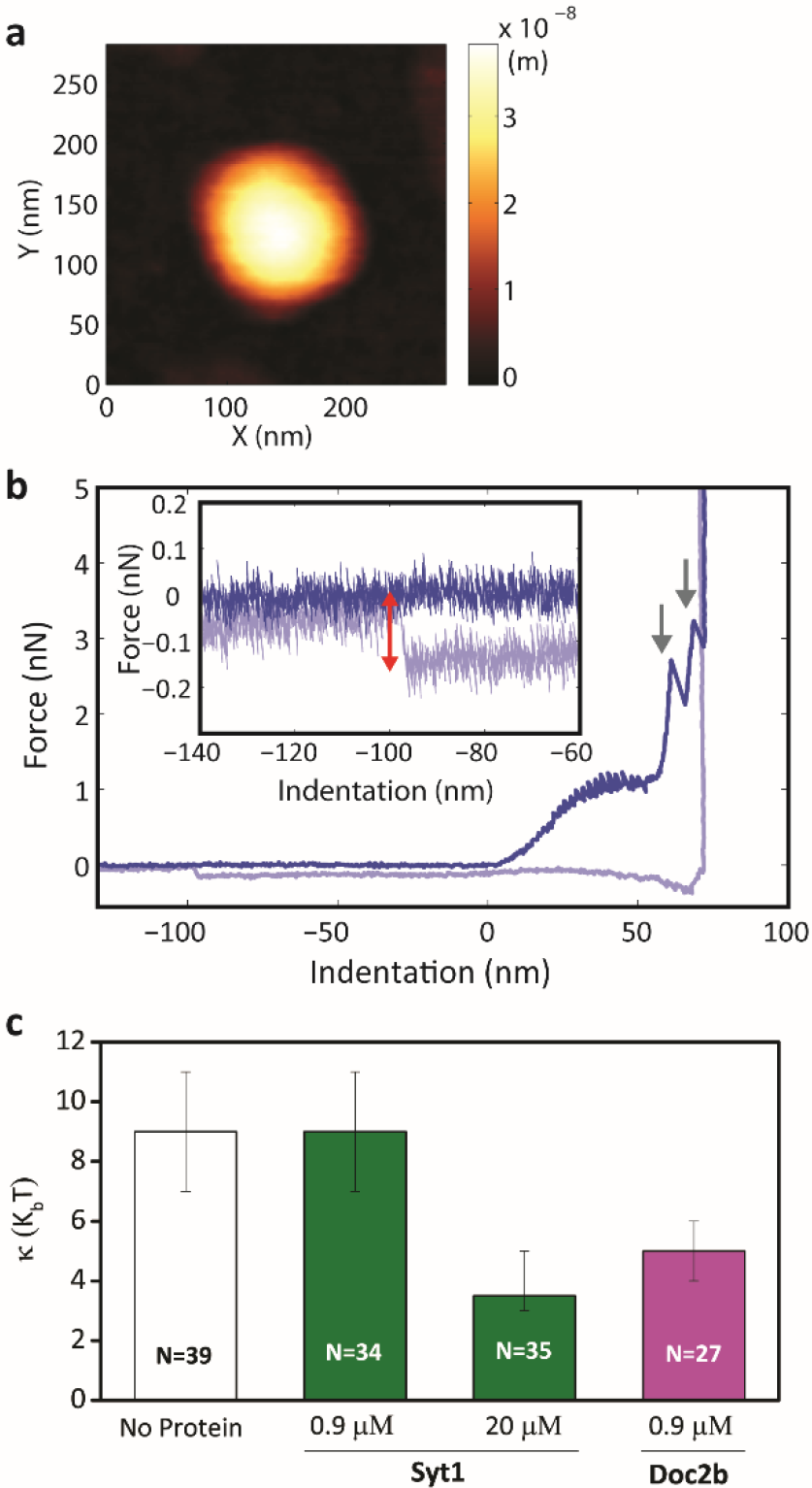
High concentrations (20 μM) of Syt1-C_2_AB and lower concentrations (0.9 μM) of Doc2b-C_2_AB reduce the membrane bending modulus. (a) AFM image of a typical vesicle, 44% cholesterol, 20% porcine brain PS, 21% egg PC and 15% Egg SM. (b) Typical force plot obtained by nanoindentation of a vesicle. From the slope of the initial linear part, the stiffness of the vesicle is obtained. Two rupture events of the two bilayers can be observed, and a tether is formed upon tip retraction. Dark blue: approach, light blue: retraction. Inset: zoom in on the tether rupture force, which is used to calculate the osmotic pressure in the vesicle as described in Vorselen et al. (36). Red arrow shows the difference in force. (c) Bending modulus values for various concentrations of Syt1-C_2_AB and Doc2b-C_2_AB, with N values indicating the numbers of indented vesicles. Error bars mark 68% confidence intervals determined by bootstrapping.

## Discussion

In this work, we observed that the C_2_AB domain of Syt1 induces the formation of membrane tethers in a Ca^2+^ and protein concentration-dependent manner. Similar to Doc2b induced membrane tethers(33), Syt1 induced tethers contained both protein (shown by Syt1-C_2_AB-mCherry labeling) and phospholipid (shown by rhodamine-PE labeling) and resisted high forces during bead separation.

Our experiments highlight several important differences between Syt1- and Doc2b-induced membrane tethers. When comparing symmetrical (both sides) and asymmetrical (one side) presence of protein on the membranes, Syt1 favors an asymmetrical and Doc2b a symmetrical configuration. This difference is reminiscent of the physiological configuration of the two proteins: Syt1 is anchored to the vesicle membrane by a transmembrane domain, and Doc2b is a soluble protein that can diffuse in the cytoplasm and has access to both the vesicle and target membranes; When expressed as a Doc2b-EGFP fusion, the fluorescence is usually homogeneously distributed in the cytosol and accumulates Ca^2+^-dependently with the plasma membrane. Using subcellular fractionation & western blotting, Doc2b was also observed in the synaptic vesicle fraction ((40), figure 3 therein, though a direct comparison between the Doc2b distribution *in vivo* and the membrane-bound configuration in our experimental setup is complicated by differences in the free Ca^2+^ concentration, membrane compositions and absence of the Doc2b N-terminal domain which contains a Munc13-binding site). For Syt1, two hypotheses pertaining to the mechanism of membrane bridging have been proposed (reviewed by Seven et al. (13)): the direct bridging hypothesis, which states that C_2_AB molecules bind simultaneously to both membranes, and the oligomerization hypothesis, in which interactions between C_2_AB oligomers formed on both membranes bridge the membrane gap. The higher activity of Syt1 in the asymmetric setup, which favors protein-membrane interactions, provides support for the direct bridging hypothesis. This notion was confirmed by the lack of Syt1-mCherry redistribution to the unlabeled bead in our experiments. Doc2b, on the other hand, was more efficient in the symmetric conformation, which may suggest a preference for protein-protein (and not protein-membrane) interactions.

Another difference is observed in membrane bridging vs. hemifusion. In confocal fluorescence microscopy of bead pairs, the ability of the fluorescent phospholipid tracer Rhodamine-PE to diffuse to an unlabeled bead was used to discriminate between two possible membrane configurations: structures without membrane continuity (likely resulting from membrane bridging), and with membrane continuity (likely representing hemi(fused) membranes as observed previously)(13). For low concentration of Syt1-C_2_AB, all 58 membrane tethers examined in our study appeared to originate from membrane bridging. This result is consistent with previous studies that demonstrated membrane bridging by Syt1 (12, 13) at similar concentrations. At 20 μM protein concentration Syt1 induced hemifusion, in the presence of cholesterol and only in the asymmetrical configuration. In contrast, Doc2b induced membrane tethers caused both membrane-bridged and hemifused structures, already at low protein concentrations.

When comparing tether rupture forces, several factors need to be taken into account. The rupture is expected to occur at the weakest point, therefore most likely at the contact between the membranes. The contact point can either be composed of membrane-bound protein (bridging), a continuous outer membrane leaflet (hemifusion), or a fully fused membrane (which has been excluded in content mixing experiments(33)). For Doc2b and Syt-1 generated membrane tethers, the rupture force was increased in the presence of 30% cholesterol, a lipid that has been suggested to assist membrane fusion(34). Due to its effective negative curvature, cholesterol has been proposed to lower the energy for forming lipid hemifusion stalks that are thought to be intermediates in membrane fusion (41, 42). As for Doc2b, the dramatic increase in rupture force beyond the measurement limit correlates with the increase in hemifusion observed in our confocal fluorescence measurements, we hypothesize that upon hemifusion, stronger tethers will be formed. Possibly, such strong tethers are not ruptured in our experiments, as the maximal forces we can apply are limited by the optical trapping power. For Syt1-C_2_AB induced tethers, the median rupture force most likely represents protein – membrane interactions with a median rupture force in the 50 – 200 pN range, strongly dependent on the Ca^2+^ (which regulates C_2_AB-membrane binding) and protein concentration. The binding force of one Syt1 molecule to a membrane is in the range of several pN(43). Therefore, the differences we observe in the force probably relate to a variation in the number of bound molecules.

Membrane fusion is energetically unfavorable and does not occur spontaneously *in vivo* (44). One major energy barrier is associated with the strong repulsive hydration forces between bilayers. Other energy-costly intermediates include lipid splaying during initiation of hemifusion-stalk formation, stalk expansion- which yields the hemifusion diaphragm, and, finally, fusion pore formation(45). In the case of synaptic vesicle fusion, the activation energy of bilayer-bilayer fusion is very high (≈40 k_B_T) and the process is aided by important proteins including SNAREs, complexin and Ca^2+^ sensors. It has been suggested that in response to Ca^2+^ binding, Syt1 could promote SNARE-mediated fusion by lowering this energy barrier via induction of positive curvature in target membranes upon insertion of C_2_-domains into the membrane(23). Using AFM, we have directly measured a significant reduction in the bending modulus of a lipid membrane upon Syt1-C_2_AB binding (Fig. 6) in the presence of 20 μM protein. For Doc2b, a significant reduction of the bending modulus was observed already at 0.9 μM protein concentration. This difference could be due to larger effective curvature for Doc2b that is expected to decrease membrane thickness (46), thus affecting the bending modulus which depends on membrane thickness to the power of three (47). A membrane inclusion with higher curvature is theoretically predicted to result in a larger reduction in the bending modulus (48). How does this global change in the mechanical properties of the membrane contribute to the local event of hemi(fusion)? A possible explanation is that membrane “dimples” form locally due to protein insertion. The curvature induction hypothesis predicts the formation of a buckle-like membrane structure between protein insertions in the membrane. Locally, the end cap membrane is highly curved and thus the lipids are under stress. This stress is partially relieved during lipid rearrangements accompanying the fusion process, which reduces the overall energy cost of the reaction, thus facilitating membrane fusion (22). We note that is difficult to compare results of bending moduli obtained by different methods, as a large variation exists in bending modulus values for the same bilayer composition obtained by different measurement techniques (49). Nonetheless, our results are in the same range as other reported values (50, 51).

Combining all observations, it is likely that membrane insertion of the C_2_AB domain of Doc2b results into a larger disruption of the lipid packing compared with Syt1, consequently reducing the energy barrier for hemifusion, and allowing hemifusion already at low protein concentrations. Syt1, however, might not sufficiently disrupt membrane structure at low concentration, and changing the configuration to asymmetrical or addition of cholesterol is not sufficient to push the membranes towards hemifusion. Only at higher concentration enough disruption to membrane packing is attained, which is manifested in lower bending modulus and concomitantly, high hemifusion probability in the asymmetrical configuration. The lowering of the energy required for membrane deformation likely contributes to the overall Ca^2+^-secretion triggering mechanism by Syt1 and Doc2b, and this mechanism may also be relevant for other C_2_AB containing Ca^2+^ sensor proteins.

## Acknowledgments

We are grateful to Michael Kozlov for helpful discussions. We further thank Ineke Brouwer for useful discussions and technical advice. RS acknowledges support through HFSP postdoctoral fellowship LT000419/2015, as well as support through the Israeli National Postdoctoral Award for Advancing Women in Science, and the L`Oreal UNESCO award for advancing women in science. WHR acknowledges the support of the Nederlandse Organisatie voor Wetenschappelijk Onderzoek Vidi grant. JR acknowledges the support of the Welch Foundation (grant I-1304) and the National Institutes of Health (Research Project Award R35 NS097333).

## Materials and methods

### Bead coating

1,2-Dioleoyl-sn-glycero-3-phosphocholine (DOPC) and 1,2-dioleoyl-sn-glycero-3-phospho-L-serine (DOPS) chlorophorm solutions were purchased from Avanti Polar Lipids. Lissamine-Rhodamine B 1,2-dihexadecanoyl-sn-glycero-3-phosphoethanolamine was purchased from Invitrogen. To prepare membrane-coated microspheres, polystyrene nonporous beads of diameter 3.84 μm±4% were acquired from Spherotech. Prior to the coating procedure, the beads were washed three times in milliQ and collected after each wash by centrifugation for 3 minutes at 900 × *g*. Liposomes were prepared by mixing lipids in chloroform solutions, extensively drying the mix under a nitrogen gas stream, and hydrating the lipid cake in milliQ to a final lipid concentration of 1 mg /ml. The suspension was then vortexed, sonicated on ice water and centrifuged for 90 min at 21,000 × *g* at 4°C. The supernatant was mixed with bead and incubated for 16 h at 4°C in presence of 3 mM CaCl_2_ with gentle continuous rotation to keep the beads dispersed. The beads were washed (each time collecting the beads for 3 minutes at 900 × *g* and gently resuspending them) in buffer 1 containing 25 mM HEPES, pH 7.4, 200 mM NaCl, 1 mM Tris 2-carboxyethyl-phosphine (TCEP) and 5 mM EDTA, then buffer 2 (25 mM HEPES, pH 7.4, 100 mM NaCl, 1 mM TCEP, 0.25 mM CaCl_2_) and then twice in buffer 3 (25 mM HEPES, pH 7.4, 25 mM NaCl, 1 mM TCEP and 0.25 mM CaCl_2_). The coated beads were stored at 4°C prior to use. To visualize lipid tethers that formed in the presence of Syt-1-C_2_AB or Doc2b-C_2_AB, we prepared coated beads containing 1% Rhodamine PE.

### Recombinant proteins

Recombinant Doc2b-C_2_AB (rat, amino-acid residues 115–412) was prepared as previously described(33). Recombinant Syt1-C_2_AB (rat, amino acid residues 140-421) or Syt1-C_2_AB-mCherry (same fragment) were cloned into the pGEX4T3 vector and transformed into the *E. coli* BL21 strain for expression. Cultures were grown in LB medium (VWR Life Science) enriched with 100 μg/mL ampicillin (Fisher bioreagents), or in LB medium supplemented with both 100 μg/mL ampicillin. Gene expression was induced with 0.1 mM IPTG (Fisher bioreagents), after which cultures were grown O/N at 34°C and 210 rpm to maximize aeration. Lysate buffer (300 mM NaCl, 50 mM Tris, pH 7.5, 10 mM EDTA, 1 mg/mL lysozyme, protease inhibitor cocktail) was added to the bacterial pellet, before the lysate was sonicated 4 times for 15 seconds with 1 minute intervals, while cooling on ice water. Pellet was incubated for 2 hours with 1% triton-X100, and then centrifuged for 30 minutes at 8500 g to dispose of any insoluble bacterial lysate. Protein purification took place by means of glutathione agarose beads (Sigma) affinity assay. Glutathione agarose beads slurry was added to the lysate and incubated O/N at 4°C and 5 rpm. Afterwards, beads were washed with high salt buffer (300 mM NaCl, 50 mM Tris, pH 7.4, 10 mM EDTA) and loaded into a disposable column. After a wash with 150 mM NaCl, 50 mM Tris, pH 7.4, 1.4 mM MgCl_2_, the column was incubated with DNAse I (Roche Diagnostics; 50U/mL) and RNAse A (Invitrogen; 50U/mL) for 15 minutes at RT. Column was washed with Low salt buffer (150 mM NaCl, 50 mM Tris, pH 7.4). Protein was cleaved from the glutathione agarose beads by incubation with thrombin (Serva) at 4°C. Protein concentration was determined with SDS PAGE stained with SYPRO ruby, using known quantities of bovine serum albumin for calibration. SYPRO ruby signal was visualized with a scan performed by the Fuji5000. For key experiments, Syt1-C_2_AB was ^15^N-labeled by expressing the protein in minimal medium containing ^15^NH_4_Cl as the sole nitrogen source, and the protein was purified by gel filtration and ion exchange chromatography as described (11). ^1^H-^15^N HSQC spectra of the protein were then used to verify the absence of polyacidic contaminants that are difficult to remove from Syt1 fragments containing the C2B domain(52, 53) These key experiments included the AFM measurements and the comparison between symmetric and asymmetric protein binding to membrane-coated beads. Proteins were aliquoted and stored at −80°C until use.

### Optical tweezers

We use a C-trap confocal fluorescence optical tweezers setup (LUMICKS) made of an inverted microscope based on a water-immersion objective (NA 1.27), together with a condenser top lens placed above the flow cell. The optical traps are generated by splitting a 1064-nm laser (10-W CW fiber laser) into two orthogonally polarized independently steerable optical traps. To steer the two traps both a one coarse-positioning piezo stepper mirror and one accurate piezo mirror were used. Optical traps were used to capture lipids-coated beads. The displacement of the trapped beads from the center of the trap was measured and converted into a force signal by back-focal plane interferometry of the condenser lens using two position sensitive detectors. The beads distance was determined by using template-matching on the real-time imaging of a bright-field movie of the trapped beads. The samples were illuminated by a bright field 875-nm LED and imaged in transmission onto a metal-oxide semiconductor (CMOS) camera.

### Confocal fluorescence microscopy

A single, pulsed laser system (ALP-745-710-SC, 20 MHz, 100-ps pulses, Fianium) was used for confocal fluorescence excitation at a wavelength of 543 nm, selected from a supercontinuum spectrum by using an AOTF (AOTFnc-VIS-TN, AA Opto-Electronic). For scanning, a fast tip/tilt piezo mirror (S-334.1SD, Physik Instrumente GmbH & Co.,max. scan rate 200 Hz) was used. For confocal detection, the emitted fluorescence was descanned, separated from the excitation by a dichroic mirror (F33-554, AHF Analysentechnik) and filtered using an emission filter (F47-586, AHF Analysentechnik). Photons were counted using fiber-coupled APDs (APDs SPCM-AQRH-14-FC, fibers SPCM-QC9, Perkin Elmer). The multimode fibers (62 μm diameter) serve as pinholes that provide background rejection. Microfluidics: a 5-channel laminar flow cell (LUMICKS) was assembled onto an automated XY-stage (MS-2000, Applied Scientific Instrumentation). The latter allowed the controlled transfer, in a highly efficient manner, of the optically-trapped beads through the channels of the flow cell, containing the proteins of interest.

### Approach-and-separation routine

Repeated approach and separation of lipid coated beads was performed as previously described(33). Briefly, one bead was kept stationary (the left trap, Fig. 1a) and the other bead was moved towards and away from it (the x dimension). We recorded the forces in x and y directions on both beads, and used the force data in the x direction of the stationary bead, unless stated otherwise. At the initial separation of ~7 microns between the beads, the force on the stationary bead is zero. During an experiment, the beads were brought into contact such that a repulsive force of ~10 pN was exerted on the stationary bead, (step with positive force in Fig. 1a), then separated. The contact time in such experiments was five seconds. If specific protein dependent interactions occur, rupture forces are recorded upon separation, observed as negative force peaks (arrows in Fig. 1a indicate rupture events). This procedure was repeated 20 times, or until a bead escaped the trap due to very strong interactions upon pulling. The speed at which the bead is retracted is known to influence rupture forces. Therefore, a constant trap speed of 2 mm s 1 was used in all experiments.

### Symmetric and Asymmetric experiments

To test the effect of protein coating configuration on interaction strength and probability, experiments were performed as follows: beads were flushed into channel 1 containing buffer 3 without protein. After catching two beads, they were moved into the protein containing channel 4. Following incubation in the protein channel for ten seconds, the beads were moved into channel 3 which contained the same buffer without protein. Symmetrical measurements were then conducted directly in channel 3. For asymmetrical measurements, one of the beads was dropped and a different non protein coated bead was caught in channel 1. Then the beads were moved into the buffer-containing channel 3 and measurements were performed. The fluorescently labelled bead was protein coated in 50% of the cases, and the presence of the protein coating on either the labelled or unlabeled membrane did not have any effect on the results.

### Data acquisition and analysis

Data analysis was performed using custom written software in python (Gitlab link: https://gitlab.com/sorkin.raya/membrane-interactions-tweezers) to extract force data at a frequency of 100 Hz. To discriminate protein mediated specific interactions from unspecific adhesion between the membranes, a force threshold of 25 pN was used throughout the analysis. Median rupture forces as presented in the figures were determined by bootstrapping with 1000 iterations, resampling 90% of the data. Error bars are standard deviation of the bootstrapped values.

### Liposome preparation for AFM

All lipid-chloroform solutions and cholesterol were purchased from Avanti Polar Lipids. First, a round bottom flask was cleaned with 96% ethanol, then washed two times with soap followed by rinsing two times with acetone. After that the round bottom flask was dried under argon flow for ~15 min. The lipids were mixed in the round bottom flask (for spectrophotometer experiments: 50:20:30 (mole-%) DOPC: DOPS: chol; for AFM experiments: 44% cholesterol, 20% porcine brain PS, 21% egg PC and 15% egg SM). The solvent was evaporated in a rotary evaporator at 100 mbar for 60 min immersed in a water bath (40 °C). After that, buffer was added to the dried lipid cake to attain a final lipid concentration of 1 mg/ml. Vortexing this mixture together with glass beads (4 mm diameter) supported the resuspension of the lipids. Subsequently, the re-suspended lipids underwent 5 freeze and thaw cycles (~20 s in liquid nitrogen followed by 5 min in a water bath at 40 °C, each) before the liposomes were extruded 31 times using a mini extruder from Avanti^®^ Polar Lipids and a 0.1 μm filter pore size. Prior to extrusion the mini extruder and syringes were thoroughly cleaned by rinsing with water and 70% ethanol. The liposomes were stored in the fridge over night and on ice throughout the measurements and used within 5 days.

### Liposome aggregation assay and analysis

The liposome aggregation assay was described in detail elsewhere(8). The spectrophotometer measurements were conducted with a Varian Cary^®^ 50 UV-Vis Spectrophotometer from Agilent Technologies. For all measurements, a wavelength of 350 nm was used, the average reading was 0.1 s and a cycle was set to 1.00 s. A precision cell (cuvette) made of Quartz SUPRASIL^®^ (light path 10.00 mm) from Hellma^®^ was used to perform the experiments. The blank measurement was done for 500 s with 170 μl buffer 3 + 40 μl liposomes (DOPC:DOPS:Chol 50:20:30 (mole-%) at 1 mg/ml). Right before each measurement the cuvette was rinsed three times with buffer 3 and then loaded with 160 μl buffer 3 and 40 μl liposomes. The liposome dispersion was gently mixed twice by flipping the cuvette over. ~87 s after the starting time of the measurement, the cuvette was taken out of the spectrophotometer. At ~100 s of the measurement 10 μl protein (titrated before) was added to one side of the precision cell and mixed twice by flipping the cuvette carefully to the injection side. Right after, the cuvette was inserted back into the spectrophotometer and the absorbance measurement was continued for a total time of 500 s. Before switching samples, the cuvette was cleaned by rinsing multiple times with buffer 3 and milliQ water and dried under a nitrogen flow.

All curves were baseline corrected: the average absorption before extracting the cuvette (first ~87 s) was calculated and subsequently subtracted from all absorbance data points of a measurement. After that, the average maximum absorbance (maximum liposome binding) and the corresponding S.E.M. values were determined for each measurement condition, e.g. each protein concentration. At least three measurements per condition were performed.

### AFM experiments

Liposomes were adhered to poly-L-lysine coated glass slides, prepared as follows: Slides were cleaned in a 96% ethanol, 3% HCl solution for 10 minutes. Next, they were coated for 1 hour in poly-L-lysine (a 0.001%, Sigma) solution, rinsed with ultrapure water, and dried 20 hr at 37 °C. They were stored at 7 °C for maximally one month. A 10 μL drop of liposomes diluted in buffer 3 was incubated on the glass slide. Vesicles were imaged in PeakForce Tapping™ mode on a Bruker Bioscope catalyst setup. Imaging was performed at RT, 22°C. Force set point during imaging was 100 pN - 200 pN. Nano-indentations were performed by first making an image of a single particle, then indenting it until a trigger force of 0.5 nN is reached, and subsequently applying higher forces (2-10 nN) at a velocity of 250 nms^−1^. Importantly, both before and after the vesicle indentation, the tip was checked for adherent lipid bilayers by recording a force-distance plot on the glass surface until a trigger-force of 5 nN. Silicon nitride tips with a nominal tip radius of 15 nm on a 0.1 N/m cantilever by Olympus (OMCL-RC800PSA) were used. Individual cantilevers were calibrated using thermal tuning.

### AFM image analysis

Both images and force curves were processed using home-built MATLAB software. Size and shape were analyzed from line profiles through the maximum of the vesicle along the slow scanning axis. Circular arcs were fitted to the part of the vesicle above half of the maximum height to obtain the radius of curvature, from which the tip radius (2 nm/ 15nm, as provided by the manufacturer) was subtracted. The height of vesicles was derived from FDCs, and the difference between the height obtained from FDCs and images was used for a subsequent correction of *R*_*c*_ (36). *R*_*0*_, the unperturbed vesicle radius in dispersion, was calculated under the assumption of surface area conservation as previously described (36).

### AFM FDC analysis

Analysis was done as described in detail previously (36). Briefly, raw data of a force cycle, given by the deflection of the cantilever versus the Z-piezo displacement, was converted to force versus separation (between the tip and the sample, or FDC) by subtracting the cantilever deflection. Contact point between tip and vesicle was found by using a change point algorithm and occasionally manually adjusted. Stiffness of the EVs was found by fitting a straight line in the interval between 0.02 – 0.1 *R*_*c*_. For finding the tether force, a step fitting algorithm based on the change point analysis, which divides the curve into segments with slope 0. Only adhesion events extending beyond the contact point were included. For fitting to the theory, described in detail elsewhere (36), the sum of the squared log Euclidian distance between the theoretical curve and the individual experimental data points was then minimized by adjusting κ as a single fitting parameter. Confidence intervals were estimated using the bias corrected percentile method with 1000 bootstrapping repetitions, for which a set of observed value combinations equal in size to the original data set was randomly drawn and fitted.

