## Supplementary material for "Synaptotagmin-1 and Doc2b exhibit distinct membrane remodeling mechanisms"


**This PDF file includes:**

Supplementary text
Figs. S1 to S5
References for SI reference citations

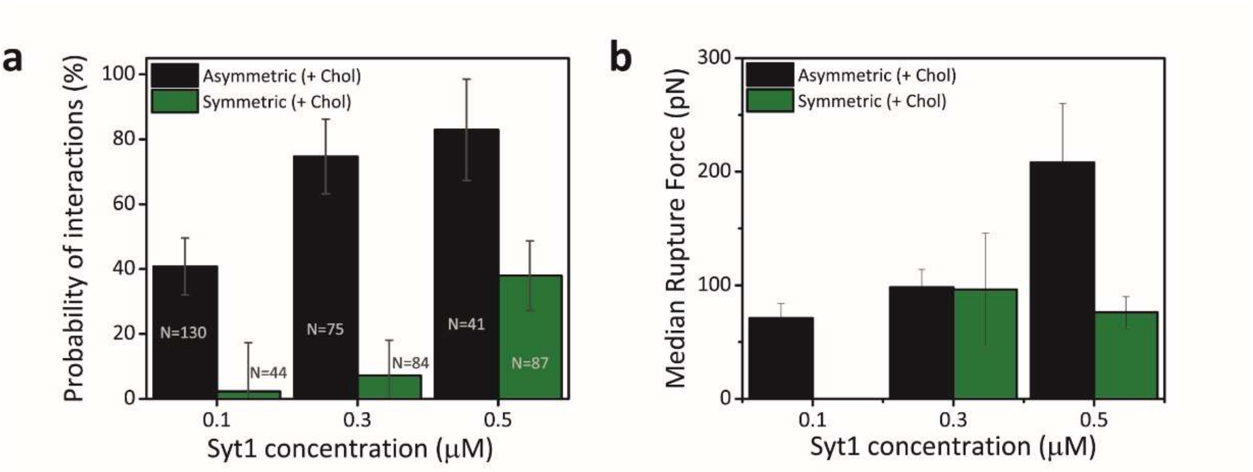


**Figure S1**: Effect of protein concentration. (a) Probability of interactions in symmetric and asymmetric configurations for different Syt1 $C_2AB$ concentrations (at 0.25 mM $CaCl_2$). For both symmetric and asymmetric configuration, the probability of interactions increases with the increase in protein concentration. (b) Median rupture forces in symmetric and asymmetric configurations for different Syt1 $C_2AB$ concentrations. Only cholesterol containing membranes were used for these measurements.

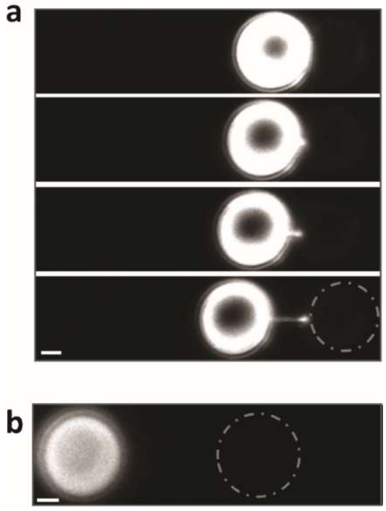


**Figure S2:** (a) Additional fluorescence images showing tether formation between an unlabelled bead (coated with PC:PS:Chol, 50:20:30) and a labelled protein-coated bead (PC:PS:Chol, 50:20:30 and 0.5 µM Syt1-$C_2$AB-mCherry), asymmetric configuration. Images were acquired while the right bead was gradually pulled away from the left bead, extending the tether that is clearly visible as it is coated by fluorescently labelled Syt1 $C_2$AB. The right bead remains dark throughout the experiment as can be seen in the images. Time between first and last image was five minutes (b) Example of a dark tether being pulled out from the right bead (coated with PC:PS:Chol, 50:20:30)while the left bead is coated with additional 1% Rhodamine-PE lipids, and dark protein (0.5 µM Syt1-$C_2$AB) is present on both beads (symmetric configuration). Scale bar: 1µm.

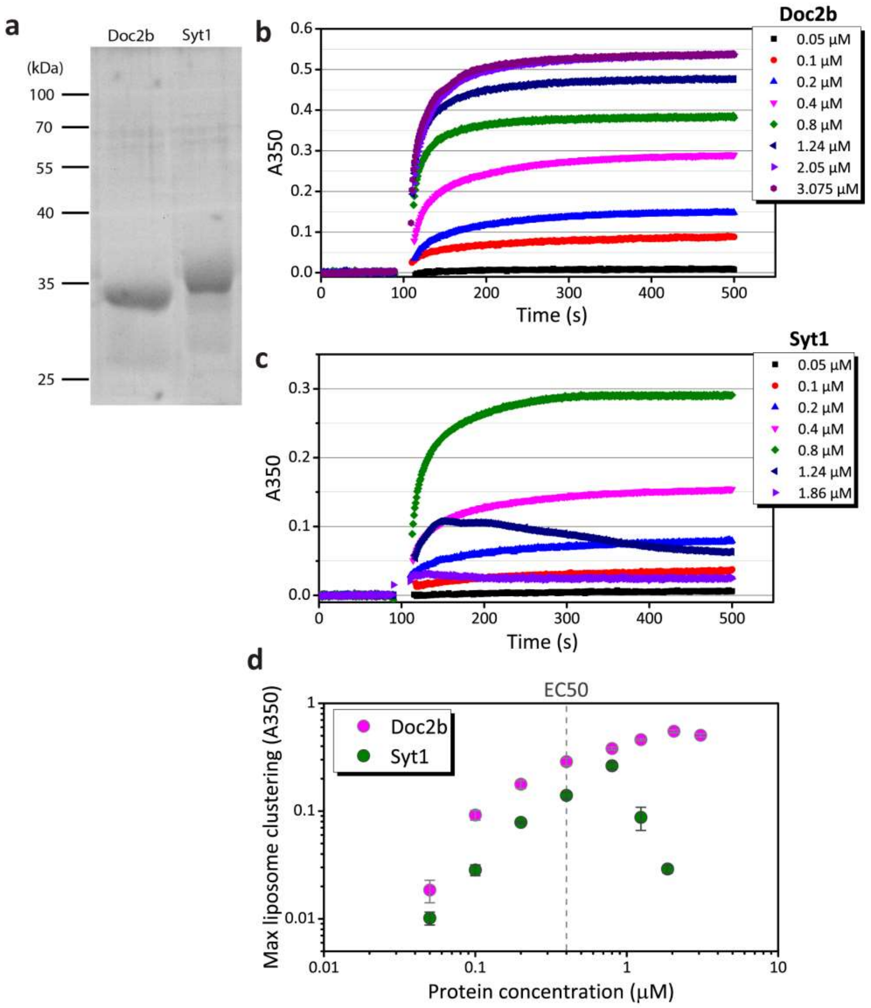


**Figure S3: (a)** SYPRO Ruby staining of recombinant Doc2b-$C_2$AB (aa.115–412) and Syt1-$C_2$AB (aa140-421) fragments separated by SDS-PAGE electrophoresis. Molecular size markers (Prestained PageRuler™) are indicated on the left. **(b)** Typical liposome aggregation measurements performed with different concentrations of Doc2b. First, the absorbance of liposomes diluted with buffer 3 were measured until t~100 s. Then, 10 µl protein at the corresponding concentration was added and carefully mixed before the cuvette was inserted again into the measurement device. The absorbance (which correlates with liposome clustering) was measured for total time of 500 s at a wavelength of 350 nm. All measurements were baseline-corrected. With increasing protein concentration an increase in the liposome clustering is observed. **(c)** Similar experiments as in (b) with Syt1-$C_2$AB. For higher protein concentrations a decrease of the liposome clustering was observed. **(d)** Maximal absorption (liposome clustering) versus protein concentration for Doc2b and Syt1. The values are averages of at least three measurements per condition. The error bars represent the standard error of the mean. While Doc2b shows an exponential behavior, Syt1 shows a more linear behavior until a maximum value is reached after which the liposome binding decreases linearly. The EC50 values for Doc2b and Syt1 are ~0.4 µM.

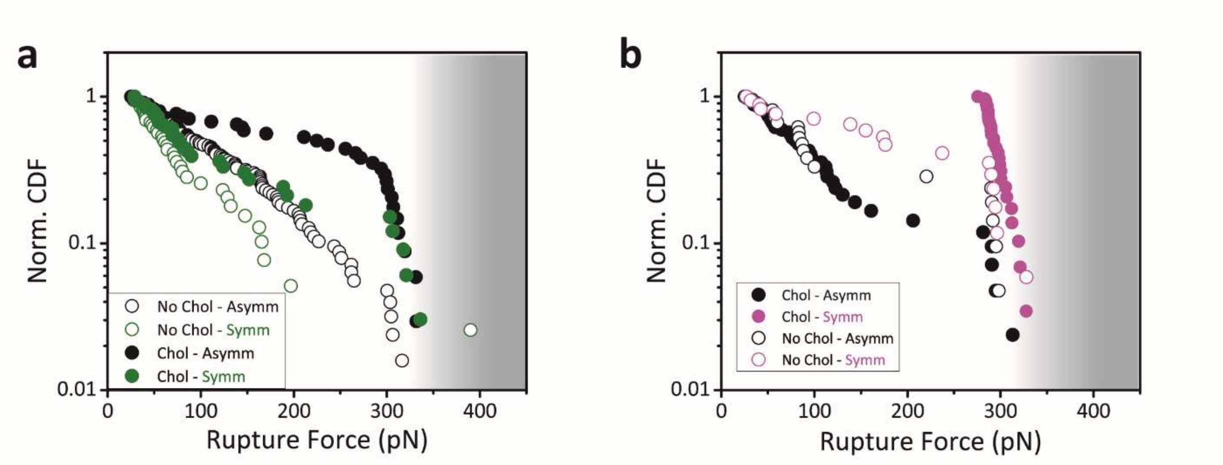


**Figure S4**: Normalized cumulative distribution functions (CDF) of the rupture forces for (a) Syt1 and (b) Doc2b, in symmetric and asymmetric configurations for each protein, with and without cholesterol in the membranes. Gradient grey background marks the maximum force region determined by the stiffness of the optical trap at the set laser power and bead size.

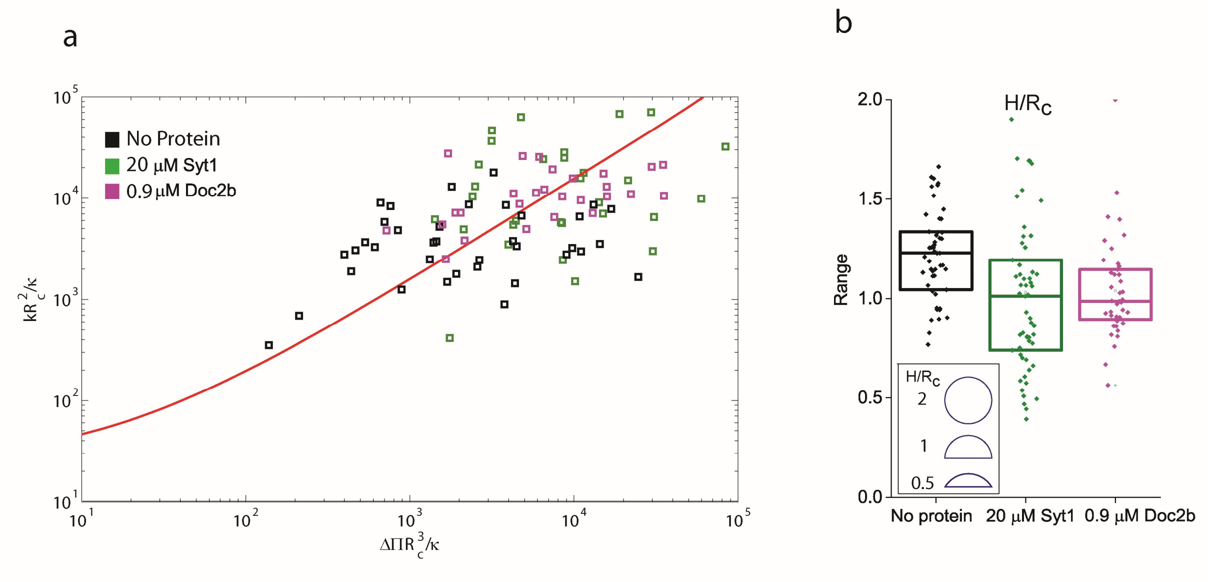


**Figure S5**: (a) Dimensionless pressure versus dimensionless stiffness for SUVs with and without added Syt1 $C_2AB$. Red line is the theoretically predicted curve. Different colored symbols are the experimental data for the two vesicle types, as indicated in the inset. Bending modulus was used as single fitting parameter. (b) Height/ radius ratio of the two vesicle populations, with and without added protein. In the presence of protein, vesicles are more flattened on the surface as they are softer.
